# *Escherichia coli* S2P family intramembrane protease RseP is engaged in the regulated sequential cleavages of FecR in the ferric citrate signaling

**DOI:** 10.1101/2021.02.04.429786

**Authors:** Tatsuhiko Yokoyama, Tomoya Niinae, Kazuya Tsumagari, Koshi Imami, Yasushi Ishihama, Yohei Hizukuri, Yoshinori Akiyama

## Abstract

*Escherichia coli* RseP, a member of the S2P family of intramembrane proteases, is involved in the activation of the σ^E^ extracytoplasmic stress response and elimination of remnant signal peptides. However, whether RseP has additional cellular functions is unclear. In this study, we attempted to identify new RseP substrates to explore still unknown physiological roles of this protease. Our mass spectrometry-based quantitative proteomic analysis revealed that the levels of several Fec system proteins encoded by the *fecABCDE* operon (*fec* operon) were significantly decreased in an RseP-deficient strain. The Fec system is responsible for the uptake of ferric citrate, and the transcription of the *fec* operon is controlled by FecI, an alternative sigma factor, and its regulator FecR, a single-pass transmembrane protein. Assays with the *fec* operon expression reporter demonstrated that the proteolytic activity of RseP is essential for the ferric citrate-dependent upregulation of the *fec* operon. Analysis using the FecR protein and FecR-derived model proteins showed that FecR undergoes sequential processing at the membrane and that RseP participates in the last step of this sequential processing to generate the N-terminal cytoplasmic fragment of FecR that participates in the transcription of the *fec* operon with FecI. Ferric citrate signal-dependent generation of this cleavage product is the essential and sufficient role of RseP in the transcriptional activation of the *fec* operon. Our study unveiled that *E. coli* RseP performs the intramembrane proteolysis of FecR, a novel physiological role that is essential for regulating iron uptake by the ferric citrate transport system.

## Introduction

While bacterial cellular membranes act as a barrier to protect a cell from extrinsic damages caused by various xenobiotics and hazardous changes in environmental conditions, they must mediate not only selective import of nutrients and other small molecules but also transduction of signals from the external milieu to adapt to environmental changes. A variety of mechanisms exist to transmit information across the membrane. Among them, regulated intramembrane proteolysis (RIP) is a crucial mechanism conserved among all kingdoms (1, 2). In RIP, a class of membrane proteases called intramembrane proteases (IMPs) mediate transmembrane signaling through the cleavage of target membrane proteins. IMPs are unique membrane-integrated proteases in that they have their proteolytic active site located within the lipid bilayer and catalyze proteolysis in the membrane. IMPs are classified into four families: site-2 protease (S2P; zinc metallopeptidase), rhomboid protease (serine protease), presenilin/signal peptide peptidase (SPP; aspartyl protease), and Rce1 (glutamyl protease) (3, 4). IMPs cleave various substrates and thereby play diverse cellular roles including stress responses, development of Alzheimer’s disease, induction of apoptosis, maintenance of mitochondrial homeostasis, invasion of apicomplexan parasites, and quality control of membrane proteins and bacterial pathogenicity (5–8).

*Escherichia coli* RseP is one of the most well-studied members of the S2P family proteases. RseP was first identified as a key factor that regulates the σ^E^ extracytoplasmic stress response through the cleavage of a single membrane-spanning anti-σ^E^ protein, RseA (9–11). In the σ^E^ extracytoplasmic stress response, the accumulation of misfolded outer membrane proteins and lipopolysaccharide biosynthesis intermediates in the cell envelope acts as stress cues to induce the cleavage of RseA by DegS, a membrane-anchored serine protease, on the periplasmic side (site-1 cleavage). This first cleavage triggers the following RseP-catalyzed second cleavage of RseA inside the membrane (site-2 cleavage), leading to liberation from the membrane of the RseA cytoplasmic domain fragment complexed with σ^E^. Finally, degradation of the RseA cytoplasmic domain fragment by cytoplasmic proteases such as ClpXP activates σ^E^ to induce the transcription of stress genes (12, 13). The first cleavage of RseA by DegS is a prerequisite for the subsequent site-2 cleavage by RseP. Similarly, most of the other S2P family proteases are known to catalyze intramembrane proteolysis of a target protein only after preceding trimming of the substrate on the extracytoplasmic side by other protease(s). *E. coli* RseP has also been shown to eliminate remnant signal peptides generated during the membrane translocation of presecretory proteins that would contribute to the quality control of the cytoplasmic membrane (14). RseP homologs (or S2P family proteases) of many gram-negative and gram-positive bacteria are involved in various cellular processes (7). They include the production of sex pheromones in *Enterococcus faecalis* (15), sporulation in *Bacillus subtilis* (16), acid response in *Salmonella enterica* (17), mucoid conversion and alginate overproduction in *Pseudomonas aeruginosa* (18), production of cholera toxin in *Vibrio cholerae* (19), and iron acquisition in *Bordetella bronchiseptica* and *P. aeruginosa* (20, 21),

RseP spans the membrane 4 times with both of its N- and C-termini facing the periplasmic space. The residues of the first and third transmembrane segments constitute the intramembrane proteolytic active site (22, 23), and the central periplasmic region between the second and the third transmembrane segments contains tandemly-arranged two PDZ domains (PDZ tandem) (24, 25) and an amphiphilic helix that is presumably involved in the proper positioning of the PDZ tandem and a substrate (26). RseP and other S2P proteases generally cleave a single-spanning membrane protein with type II (NIN-COUT) topology. We previously proposed a model that the bulky PDZ tandem acts as a size-exclusion filter to prevent the access of a substrate with a large periplasmic domain to the protease active site in the membrane (25, 27). According to this model, the DegS-cleaved form of RseA that has lost most of its periplasmic domain (leaving a ~30 a. a. C-terminal periplasmic tail), but not the full-length RseA, can pass through the PDZ filter and access the active site of RseP. Cleavage of other known substrates (remnant signal peptides (14) and a small membrane protein, YqfG (28)) can also be explained by this model, because the formers are generated by the preceding cleavage of precursor secretory proteins around the periplasmic surface of the cytoplasmic membrane by a leader peptidase and this processing is required for the cleavage of signal peptides by RseP, whereas the latter intrinsically has a very small (~12 a. a.) periplasmic domain. The transmembrane segments of the known substrates of RseP share no detectable homology in their primary sequences, suggesting that RseP does not recognize a specific sequence motif(s) in the transmembrane segment of a substrate for its cleavage. Mutational analysis of the transmembrane segments of model substrates showed that the stability of their helical structures is an important determinant for their susceptibility to RseP (28, 29). Consistently, RseP homologs in other bacteria cleave a variety of membrane proteins with no apparent sequence homology. The sequence diversity of the RseP/S2P substrates makes it difficult to predict potential substrates from a simple sequence analysis of transmembrane sequences of membrane proteins. It would thus be reasonable to assume that RseP still has unidentified substrates and plays important roles in some cellular processes by cleaving these substrates and a more comprehensive approach is needed to search for novel substrates and cellular functions of RseP.

In this study, we employed a proteomic approach to achieve the above objective. Proteomic analysis is a powerful technique to identify novel substrates of proteases as reported previously (30–33). We found that the accumulation of several proteins encoded in the *fecABCDE* operon (*fec* operon) decreased considerably in the RseP-deficient strain. The Fec proteins constitute the Fec system that mediates the uptake of ferric citrate by a cell (34, 35). We demonstrated that FecR receives sequential processing at the membrane and that RseP plays an essential role in the activation of FecI, an alternative sigma factor dedicated to the transcription of the *fec* operon (36–39), through the regulated intramembrane cleavage of a periplasmically processed form of FecR, the regulator of the *fec* operon expression (40–42), which produces the N-terminal cytoplasmic tail fragment of FecR having a “pro-sigma” activity (40, 43) as the last step in the sequential processing of FecR at the membrane. These results uncover the details of the FecR processing in the ferric citrate-induced transmembrane signaling and a new physiological role of RseP in the iron uptake in response to the environmental conditions in *E. coli*.

## Results

### Search for new physiological substrates of RseP by quantitative proteomic analysis

For comprehensive understanding of the cellular functions of RseP in *E. coli*, we attempted to identify new physiological substrates of RseP by a quantitative proteomic approach. We expected that this analysis would enable us to find not only direct proteolytic substrates of RseP but also proteins whose expression is regulated as a result of substrate proteolysis by RseP. We compared the protein levels in the membrane fractions prepared from the Δ*rseA* Δ*rseP* mutant strain expressing the wild type or a proteolytically inactive form (a mutant having a Glu-23 to Gln alteration (E23Q) in the conserved zinc metallopeptidase active site motif, HE^23^xxH) of RseP from a plasmid. Although the *rseP* gene is essential for growth, it can be deleted in the absence of the functional *rseA* gene encoding anti-σ^E^ protein RseA, a physiological substrate of RseP. In both the strains, σ^E^ is constitutively activated because of the absence of RseA and thus we can exclude the possible changes in protein expression resulting from RseP-dependent induction of the σ^E^ stress response. The membrane fractions were prepared from these strains grown to mid-log phase in LB broth containing 1 mM IPTG, an inducer of the plasmid-encoded RseP derivatives, and subject to nanoLC/MS/MS analysis. As a result, we identified 13,815 unique peptides derived from 1,419 proteins (*E. coli* strain K-12). Among the identified proteins, 17 exhibited a significant (*P* <0.05) increase whereas 41 exhibited a significant decrease in the samples prepared from the RseP(E23Q)-expressing strain compared with the samples prepared from RseP(WT)-expressing strain (Table 1). Gene ontology enrichment analysis using the DAVID (44, 45) showed that the former included groups related to TonB box receptor and transmembrane beta strand, and the latter included proteins related to iron transport (Fold Enrichment >10, Table S1). Among the former proteins, three [CyoE (heme O synthase (46, 47)), AmtB (ammonium transporter (48)), and ZupT (heavy metal divalent cation transporter (49, 50))], exhibited a fold change greater than 2 (Fig. 1). Although these proteins accumulated under the RseP-deficient condition, they are unlikely to be the direct substrates of RseP, as S2P proteases generally cleave a single membrane-spanning protein with type II membrane topology. The top three of the latter group were FecA, FecD, and FecE, which also exhibited a fold change greater than 2. These proteins constitute the Fec system, one of the iron uptake systems in *E. coli*. Expression of these proteins is known to be induced in response to the availability of environmental iron. The decrease in their levels in the RseP-deficient strain indicates that they should also not be the direct substrates of RseP. However, because it has been shown that many bacterial S2P proteases are involved in cellular responses to environmental changes including the availability of iron (20, 21, 51), we further examined the possible involvement of RseP in the regulation of the Fec system gene expression.

**Figure 1.**
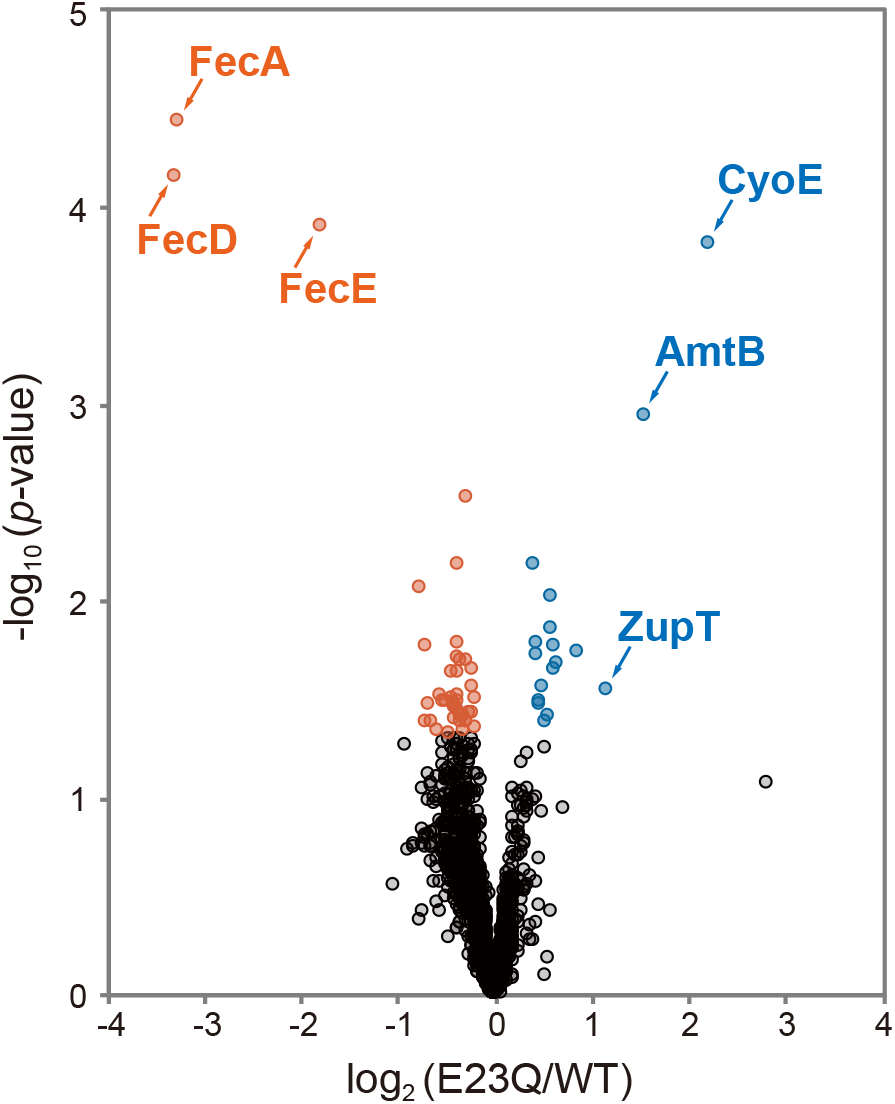
The RseP-dependent changes of the proteomic profile of the *E. coli* membrane fraction. The identified proteins were ranked in a volcano plot according to their statistical *p-*value (-log10 *p-*value, *y-*axis) and their relative abundance ratios (log2 fold-change, *x-*axis) between membrane fraction from the wild-type RseP(WT) or a protease active site mutant RseP(E23Q) expressing strains. KK377 (Δ*rseA* Δ*rseP*) cells carrying pKK47 (RseP(WT)) or pYK2 (RseP(E23Q)) were grown at 30°C in L medium containing 1 mM IPTG and 1 mM cAMP until mid-log phase. The membrane fractions were prepared by sonical disruptions of cells followed by ultracentrifugation and suspended in PTS solution. After reduction, extracted proteins were digested and the resultant peptides were subjected to nanoLC/MS/MS analyses. The proteins with *p-*value < 0.05 and log2 fold-change < 0 are indicated by red cicles, and *p-*value < 0.05 and log2 fold-change > 0 are indicated by blue circles, and the other proteins are indicated by black circles.

**Table. 1.** Significantly changed proteins (*p-*value < 0.05) depending on the proteolytic activity of RseP.

UP regulated
| Log<sub>2</sub><br>(E23Q/WT) | -Log<sub>10</sub><br>$p$-value | Gene<br>names | Protein names | Locati<br>on |
| --- | --- | --- | --- | --- |
| 2.20 | 3.80 | *cyoE* | Protoheme IX farnesyltransferase | IM |
| 1.54 | 2.94 | *amtB* | Ammonia channel | IM |
| 1.14 | 1.55 | *zupT* | Zinc transporter ZupT | M |
| 0.85 | 1.73 | *ycfJ* | Uncharacterized protein YcfJ | M |
| 0.63 | 1.67 | *fiu* | Catecholate siderophore receptor Fiu | OM |
| 0.62 | 1.65 | *hdeB* | Acid stress chaperone HdeB | Peri |
| 0.61 | 1.76 | *osmB* | Osmotically-inducible lipoprotein B | M |
| 0.58 | 2.01 | *mdtA* | Multidrug resistance protein MdtA | IM |
| 0.57 | 1.85 | *cirA* | Colicin I receptor | OM |
| 0.55 | 1.41 | *wzc* | Tyrosine-protein kinase wzc | IM |
| 0.52 | 1.38 | *glnB* | Nitrogen regulatory protein P-II 1 | - |
| 0.48 | 1.56 | *yjiY* | Inner membrane protein YjiY | IM |
| 0.46 | 1.46 | *rseP* | Regulator of sigma-E protease RseP | IM |
| 0.46 | 1.49 | *fepA* | Ferrienterobactin receptor | OM |
| 0.44 | 1.72 | *yqiC* | Uncharacterized protein YqiC | Cyto |
| 0.44 | 1.77 | *zntA* | Lead, cadmium, zinc and mercury-transporting ATPase | IM |
| 0.39 | 2.18 | *mdtB* | Multidrug resistance protein MdtB | IM |

DOWN regulated
| Log<sub>2</sub><br>(E23Q/WT) | -Log<sub>10</sub><br>$p$-value | Gene<br>names | Protein names | Locati<br>on |
| --- | --- | --- | --- | --- |
| -3.30 | 4.15 | *fecD* | Fe(3+) dicitrate transport system permease protein FecD | IM |
| -3.27 | 4.42 | *fecA* | Fe(3+) dicitrate transport protein FecA | OM |
| -1.80 | 3.90 | *fecE* | Fe(3+) dicitrate transport ATP-binding protein FecE | IM |
| -0.76 | 2.06 | *yeeR* | Inner membrane protein YeeR | IM |
| -0.72 | 1.76 | *znuA* | High-affinity zinc uptake system protein ZnuA | Peri |
| -0.71 | 1.39 | *flu* | Antigen 43; Antigen 43 alpha chain; Antigen 43 beta chain | Peri |
| -0.69 | 1.47 | *sucD* | Succinyl-CoA ligase [ADP-forming] subunit alpha | - |
| -0.64 | 1.39 | *yaaA* | UPF0246 protein YaaA | - |
| -0.58 | 1.34 | *putA* | Bifunctional protein PutA; Proline dehydrogenase; Delta-1-pyrroline-5-carboxylate dehydrogenase | - |
| -0.56 | 1.51 | *cstA* | Carbon starvation protein A | IM |
| -0.54 | 1.48 | *truB* | tRNA pseudouridine synthase B | - |
| -0.50 | 1.48 | *purN* | Phosphoribosylglycinamide formyltransferase | - |
| -0.48 | 1.32 | *rluB* | Ribosomal large subunit pseudouridine synthase B | - |
| -0.44 | 1.50 | *accA* | Acetyl-coenzyme A carboxylase carboxyl transferase subunit alpha | Cyto |
| -0.44 | 1.62 | *ycbX* | Uncharacterized protein YcbX | - |
| -0.43 | 1.39 | *thrS* | Threonine--tRNA ligase | Cyto |
| -0.43 | 1.45 | *thiI* | tRNA sulfurtransferase | Cyto |
| -0.41 | 1.48 | *gltA* | Citrate synthase | - |
| -0.41 | 1.45 | *sodB* | Superoxide dismutase [Fe] | - |
| -0.40 | 1.78 | *hrpA* | ATP-dependent RNA helicase HrpA | - |
| -0.40 | 1.48 | *rne* | Ribonuclease E | Cyto |
| -0.39 | 2.19 | *pepP* | Xaa-Pro aminopeptidase | Cyto |
| -0.39 | 1.51 | *serC* | Phosphoserine aminotransferase | Cyto |
| -0.39 | 1.71 | *ybeD* | UPF0250 protein YbeD | - |
| -0.39 | 1.64 | *selB* | Selenocysteine-specific elongation factor | Cyto |
| -0.38 | 1.44 | *degP* | Periplasmic serine endoprotease DegP | IM |
| -0.35 | 1.41 | *tehB* | Tellurite methyltransferase | Cyto |
| -0.35 | 1.38 | *gmhA* | Phosphoheptose isomerase | Cyto |
| -0.34 | 1.70 | *oppF* | Oligopeptide transport ATP-binding protein OppF | IM |
| -0.33 | 1.42 | *uvrA* | UvrABC system protein A | Cyto |
| -0.31 | 1.34 | *rlmJ* | Ribosomal RNA large subunit methyltransferase J | - |
| -0.30 | 1.38 | *clpP* | ATP-dependent Clp protease proteolytic subunit | Cyto |
| -0.30 | 2.52 | *helD* | Helicase IV | - |
| -0.29 | 1.38 | *cspB* | Cold shock-like protein CspB | Cyto |
| -0.29 | 1.69 | *fadD* | Long-chain-fatty-acid--CoA ligase | IM |
| -0.27 | 1.42 | *ybaY* | Uncharacterized lipoprotein YbaY | IM |
| -0.24 | 1.55 | *yciO* | Uncharacterized protein YciO | - |
| -0.23 | 1.64 | *lrp* | Leucine-responsive regulatory protein | - |
| -0.22 | 1.42 | *pncB* | Nicotinate phosphoribosyltransferase | - |
| -0.21 | 1.49 | *fdhE* | Protein FdhE | Cyto |
| -0.21 | 1.35 | *ahpF* | Alkyl hydroperoxide reductase subunit F | - |

### RseP function is required for the transcriptional activation of the *fecABCDE* operon

Since iron is generally an essential micronutrient for living organisms, cells now have many evolved iron-uptake systems (52). An *E. coli* cell is also equipped with multiple iron uptake systems. Among them, the Fec system acts in the uptake of ferric ion (Fe^3+^) in the form of ferric citrate (Fig. 2A). Ferric citrate in the extracellular milieu is first transported into the periplasmic space by FecA, an outer membrane transporter (53–55), and then imported into the cytoplasm via the ABC transporter FecCDE with the assistance of the periplasmic protein FecB (56). These Fec proteins are expressed from the single *fecABCDE* operon (*fec* operon) whose transcription is under the control of an alternative sigma factor, FecI (36–39). Because the genes for FecA, FecD, and FecE that exhibited decreased accumulation in the RseP-deficient strain belong to the same *fec* operon (38), we supposed that RseP is involved in the upregulation of this operon, and examined this possibility.

**Figure 2.**
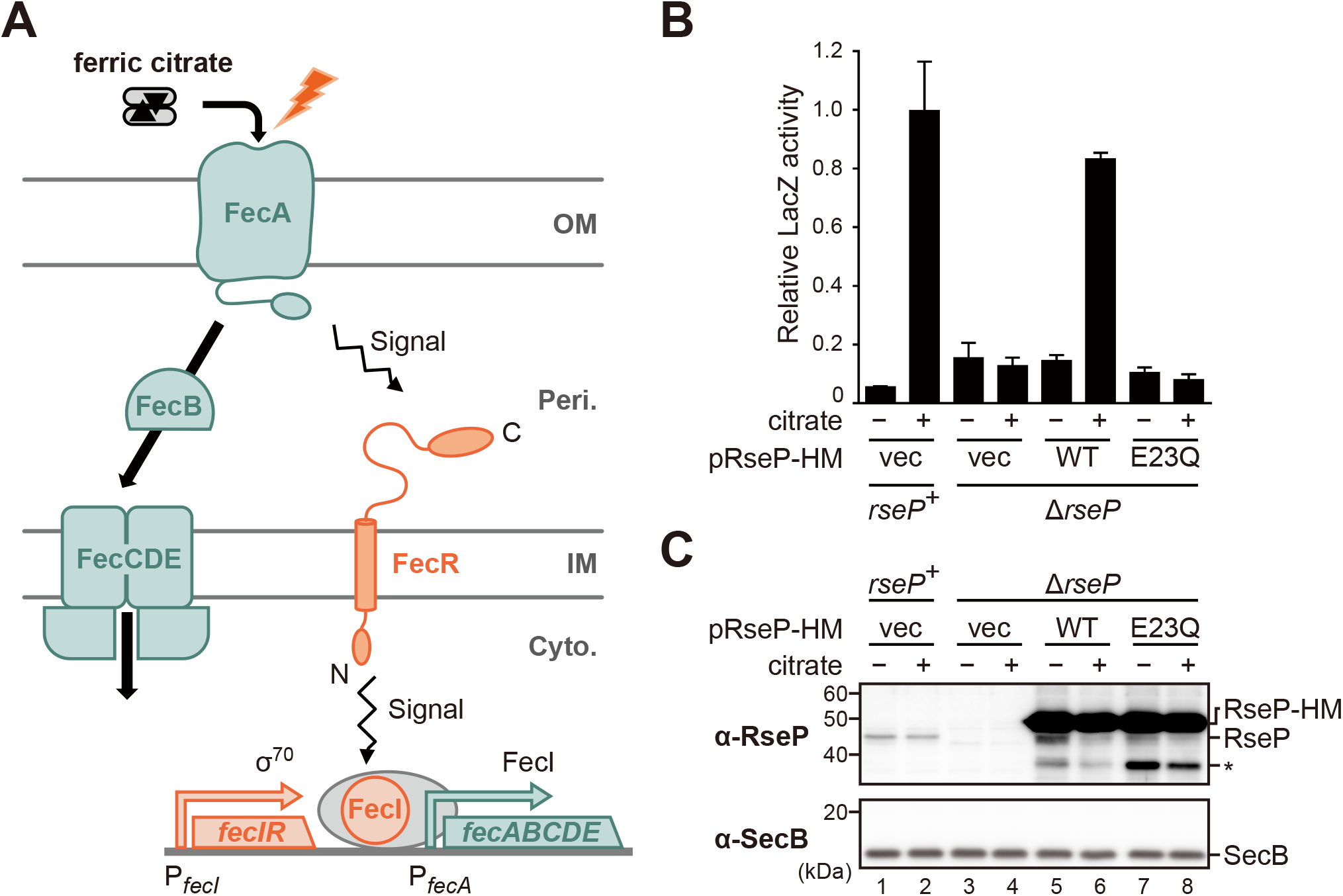
The proteolytic function of RseP is necessary for the ferric citrate-dependent transcriptional activation of the *fec* operon. (A) A schematic representation of the Fec system. Ferric citrate is transported by outer membrane transporter FecA into the periplasm and then iron is transported into the cytoplasm by periplasmic protein FecB and ABC transporter FecCDE. FecR, a type-II (Nin-Cout) single-pass inner membrane protein, receives the signal of ferric citrate from FecA and transmits it to FecI, an alternative sigma factor. Activated FecI induces the transcription of the *fecABCDE* operon (*fec* operon). The expression of the *fecIR* operon encoding FecI and FecR is independent of the FecI activity. OM, Peri, IM, and, Cyto indicate outer membrane, periplasm, inner membrane, and cytoplasm, respectively. (B) RseP dependency of the PfecA-*lacZ* reporter expression in response to ferric citrate. YH426 (Δ*ompA* Δ*ompC rseP*^+^) or KA363 (Δ*ompA* Δ*ompC* Δ*rseP*) cells harboring pYK149 (PfecA-*lacZ*) and pSTD343 (*lacI*) were further transformed with pSTD1060 (vec), pYK151 (RseP(WT)-HM, WT) or pYK153 (RseP(E23Q)-HM). The cells were inoculated into M9-based medium containing 1 mM IPTG, 1 mM cAMP, and 0.1 μM FeCl3 supplemented with or without 1 mM Na3-citrate, grown at 30°C until mid-log phase and their LacZ activities were measured. The relative LacZ activities normalized by the LacZ activity of the *rseP*^+^ cells grown in medium containing 1 mM Na3-citrate are shown (see Experimental procedures for details). Two independent experiments were carried out and the mean values are shown with standard deviations. (C) Cellular accumulation of the RseP proteins. The cells were grown as in (B) and the proteins were analyzed by 12.5% Laemmli SDS-PAGE and anti-RseP (α-RseP) or anti-SecB (α-SecB) immunoblotting. *RseP-HM* shown on the right indicates RseP(WT)-HM or RseP(E23Q)-HM expressed from pKK47 (RseP(WT)) or pYK2 (RseP(E23Q), and *RseP* indicates the chromosomally-encoded RseP. The asterisk indicates a possible degradation product of RseP(WT)-HM or RseP(E23Q)-HM. Cytoplasmic protein SecB serves as a loading control. The positions of molecular size markers (in kDa) are shown on the left. Note that the band intensity of RseP-HM is saturated.

To easily monitor the transcription of the *fec* operon, we constructed a reporter (*fec* reporter) plasmid in which the *lacZ* reporter gene was placed under the FecI-dependent promoter of the *fec* operon (PfecA). The Δ*ompA* Δ*ompC* cells (used as the *rseP*^+^ strain) carrying the reporter plasmid exhibited about 10-fold higher LacZ activity when 1 mM Na3 citrate was added to the medium containing 0.1 μM FeCl3 (Fig. 2B, *rseP*^+^). Note that the addition of increasing concentrations of FeCl3 to the medium led to a drastic decrease in LacZ activity in both Δ*ompA* Δ*ompC* (Fig. S1A) and wild-type (*ompA*^+^ *ompC*^+^) cells (Fig. S1B). This effect was presumably caused by the negative regulation of PfecA by Fur, a global transcriptional regulator for iron homeostasis (38, 39, 41, 57). These results showed that our reporter plasmid can be used to evaluate the transcription from PfecA in response to ferric citrate.

We then introduced the reporter plasmid into the Δ*ompA* Δ*ompC* Δ*rseP* cells (*rseP* can be deleted in a strain lacking the two outer membrane proteins OmpA and OmpC (58)) and examined the effect of *rseP* disruption on the transcriptional activation of the *fec* operon in response to ferric citrate. In sharp contrast to *rseP*^+^ cells, the expression of the *fec* operon as revealed by LacZ activity was not increased by the addition of citrate (Fig. 2B, Δ*rseP* / vec). Expression of the wild-type RseP with a C-terminal His6-Myc tag (hereafter RseP-HM), but not its E23Q derivative, from another plasmid restored a citrate-dependent reporter expression (Fig. 2B, WT and E23Q), while the anti-RseP immunoblotting showed that the accumulation levels of the expressed RseP proteins were comparable (Fig. 2C). These results strongly suggest that the proteolytic activity of RseP is essential for the ferric citrate-dependent transcriptional activation of the *fec* operon.

### Proteolytic function of RseP is involved in the processing of the FecR protein

The transcription of the *fec* operon is known to be controlled by FecI, an alternative sigma factor whose activity is regulated by FecR (40–42) (Fig. 2A). FecR is a single-spanning cytoplasmic membrane protein of type II topology, similar to other RseP substrates, and has been reported to be processed into several fragments *in vivo* (59, 60). In addition, its N-terminal cytoplasmic region has been suggested to exhibit an activity (“pro-sigma” activity) that is required for the function of FecI (40, 43). These facts led us to examine the possibility that FecR is cleaved by RseP to activate FecI.

We first investigated whether RseP could affect the *in vivo* processing of FecR. To this end, we constructed a derivative of FecR with an N-terminal 3xFLAG tag (F-FecR) (Fig. 3A). We also constructed a new Δ*fecR* (Δ*fecR1*) strain in which a part of the chromosomal *fecR* gene was deleted so that the deletion totally disrupts the FecR function but negligibly affects the expression of the upstream and downstream genes (see Supplementary Experimental Procedures). Assays with the *fec* reporter using the Δ*fecR* cells showed that the expression of F-FecR restored the ferric citrate-dependent induction of transcription from P*fecA* that was abolished in the Δ*fecR* cells, indicating that F-FecR is functional (Fig. 3B). We examined the effects of *rseP* deletion on the processing of the FecR protein (Fig. 3C). F-FecR was expressed in a Δ*rseA rseP*^+^ and a Δ*rseA* Δ*rseP* strain by growing the cells in M9 medium supplemented with 1 mM Na3-citrate and 10 μM FeCl3 (Fig. 3C, *lanes 9* and *10*). Note that 10 μM FeCl3 was added to the medium during the analyses of the FecR processing to obtain clear and reproducible results (see Supplementary Results). Anti-FLAG immunoblotting with total cellular proteins showed that in addition to the full-length F-FecR band (~37 kDa; FL), several smaller FecR-derived fragments were accumulated; FL and an ~25 kDa band (labeled as CL(a)) were detected in both *rseP*^+^ and Δ*rseP* strains, whereas a ~17 kDa band (CL(b)) and a ~15 kDa band (CL(c)) were detected only in *rseP*^+^ and Δ*rseP* strains, respectively. Also, CL(b) and CL(c) were generated in a citrate-dependent manner (Fig. 3C, compare *lanes 5* and *6* with *9* and *10*). While CL(a) was generated irrespective of the addition of citrate, the accumulation levels of CL(a) were considerably higher in the absence of citrate than in its presence (Fig. 3C, compare *lanes 5* and *6* with *9* and *10*). Taken together, these results suggest that ferric citrate affects the generation and/or accumulation of FecR species in the cells. All these species (CL(a), CL(b) and CL(c)) should be N-terminal fragments of F-FecR as they retained the N-terminal FLAG tag, and most likely are the degradation products of FecR. Citrate did not affect the accumulation levels of the RseP proteins in these strains. When wild-type RseP-HM was ectopically expressed in the Δ*rseP* stain, CL(b) disappeared and instead CL(c) was detected, while no such effect was observed with the expression of the protease active-site mutant RseP(E23Q) (Fig. 3C, *lanes 11* and *12*). These results suggest that CL(c) is generated by the RseP-dependent proteolytic cleavage of the FecR-related proteins.

**Figure 3.**
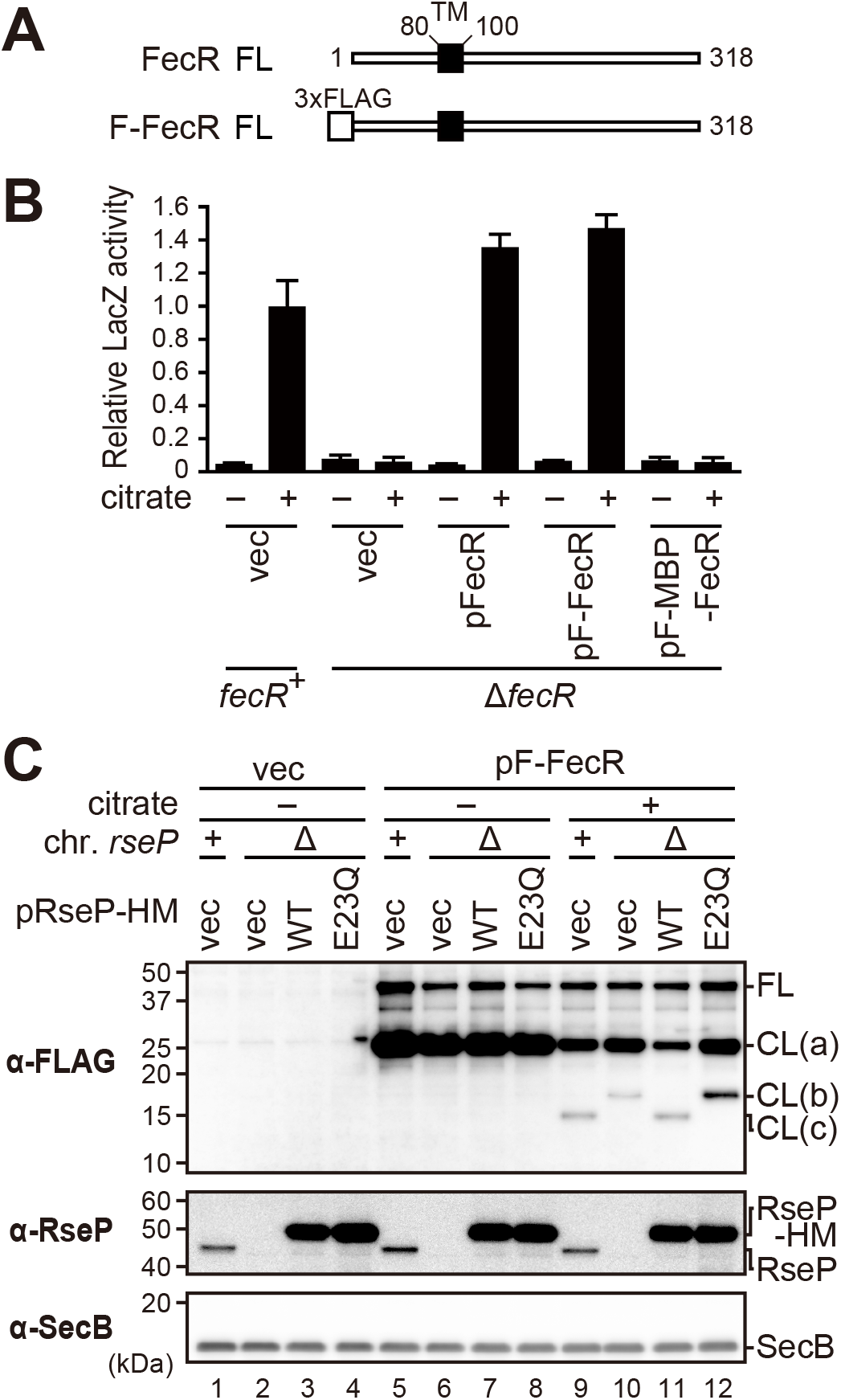
RseP is involved in the proteolytic processing of FecR. (A) Schematic representations of FecR and the N-terminally 3xFLAG-tagged FecR model substrate (F-FecR). The predicted transmembrane region and the 3xFLAG tag are shown as the black and the white boxes, respectively. *FL*, full length F-FecR. (B) Ability of various FecR model substrates to transmit the ferric citrate signal. MC4100 (*fecR*^+^) or YK627 (Δ*fecR*) cells harboring pYK149 (PfecA-*lacZ*) and pSTD343 (*lacI*) were further transformed with pSTD1060 (vec), pYK186 (pFecR), pYK188 (pF-FecR) or pYK190 (pF-MBP-FecR). The cells were grown at 30°C in M9-based medium containing 1 mM IPTG, 1 mM cAMP, and 0.1 μM FeCl3 supplemented with or without 1 mM Na3-citrate, and the LacZ activity was measured as in Fig. 2B. The relative LacZ activities normalized by the LacZ activity of *fecR*^+^ cells grown in medium containing 1 mM Na3-citrate are shown. (C) Behaviors of the F-FecR protein in the presence or absence of the co-expression of RseP and the addition of Na3-citrate. YK167 (Δ*rseA*, *rseP*^+^) or YK191 (Δ*rseA* Δ*rseP*, Δ*rseP*) cells harboring pTWV228 (vec) or pYK130 (pF-FecR) were further transformed with pSTD689 (vec), pYH9 (RseP-HM, WT) or pYH13 (RseP E23Q-HM, E23Q). The cells were grown at 30°C in M9-based medium containing 1 mM IPTG, 1 mM cAMP, 10 μM FeCl3 supplemented with or without 1 mM Na3-citrate and grown as in Fig. 2B. The proteins were analyzed by 15% Bis-Tris SDS-PAGE and anti-FLAG (α-FLAG), or 12.5% Laemmli SDS-PAGE and anti-RseP (α-RseP), or anti-SecB (α-SecB) immunoblotting. FL, CL(a), CL(b), and CL(c) on the right indicate full-length F-FecR, the possible cleavage product (a), (b), and (c), respectively. Note that the band intensities of F-FecR FL, F-FecR CL(a) and RseP-HM are saturated.

### RseP cleaves the transmembrane region of FecR to yield the cytoplasmic tail fragment

In the above immunoblotting experiments, we observed significant variations in the relative amounts of the FecR-related species (data not shown). We suspected that these variations could be caused by the instability of these species in a cell. In addition, the positive feedback regulation of the Fec system (34, 35) could complicate the results as it could enhance the ferric citrate-dependent signal and as a result the signal-induced processing of FecR. To circumvent these problems, we constructed a FecR-derived model substrate, F-MBP-FecR, in which the entire cytoplasmic domain of FecR was replaced with a tightly folded MBP (maltose binding protein) domain (Fig. 4A). This approach has proved to be useful to analyze the RseP-mediated or periplasmic proteolysis in our past studies (14, 29). The expression of F-MBP-FecR in the Δ*fecR* strain did not activate the expression of the *fec* reporter in the presence or absence of citrate, suggesting that it is not functional, as expected from its lack of the cytoplasmic domain required for the interaction with and activation of FecI (Fig. 3B). Next, we examined the *in vivo* processing of the F-MBP-FecR protein by immunoblotting (Fig. 4B and S2). In the presence of citrate, we obtained essentially the same results as those obtained with F-FecR; we detected four F-MBP-FecR-derived species that should correspond to FL, CL(a), CL(b) and CL(c), ascertaining from their sizes, and CL(c) was generated in the presence of chromosomal or ectopically expressed functional RseP, whereas CL(b) was detected instead of CL(c) under RseP-deficient conditions. Without the addition of citrate, the amounts of CL(a) were increased substantially both in the presence and absence of functional RseP, which also reproduced the results with F-FecR (Fig. 4B, compare *lanes 5*-*8* with *9*-*12*). Remarkably, even in the absence of citrate, small but significant amounts of the CL(b) and CL(c) bands were detected with F-MBP-FecR and the pattern of their generation was essentially the same as that in the presence of citrate, whereas we did not observe CL(b) and CL(c) with F-FecR in the absence of citrate. Quantification of the band intensities demonstrated high reproducibility of the results with F-MBP-FecR (Fig. S2). These observations suggest that (i) F-MBP-FecR retains its “responsibility” to ferric citrate with respect to the generation of the N-terminal fragments, although it is not functional in signal transduction and no longer enhances the Fec signaling through its positive feedback loop, and (ii) the replacement of the cytoplasmic domain of FecR with the F-MBP domain stabilizes the N-terminal fragments, which would allow for the detection of CL(b) and CL(c) without the addition of ferric citrate. These results demonstrate that F-MBP-FecR can be used as a suitable model substrate for studying of the FecR processing.

**Figure 4.**
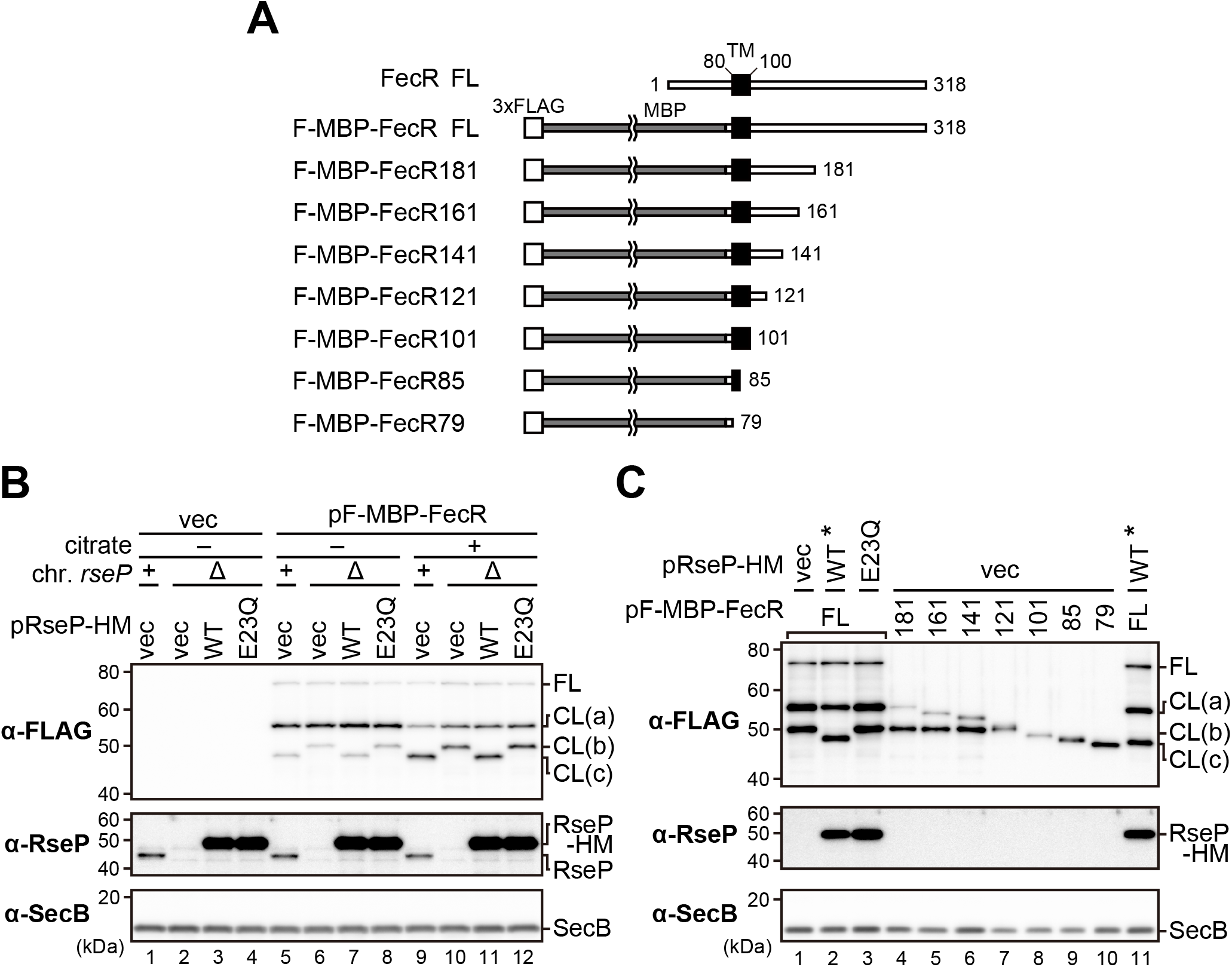
RseP cleaves the transmembrane region of FecR. (A) Schematic representations of FecR, the F-MBP-FecR model substrate, and the truncated derivatives of F-MBP-FecR. The regions corresponding to the FecR-derived cytoplasmic and periplasmic domains are shown as white lines, MBP as the gray line, the predicted transmembrane region as a black box, and 3xFLAG tag as a white box, respectively. (B) Behavior of the F-MBP-FecR protein in response to the co-expression of RseP and the addition of Na3-citrate. YK167 (Δ*rseA*, *rseP*^+^) or YK191 (Δ*rseA* Δ*rseP*) cells harboring pTWV228 (vec) or pYK147 (pF-MBP-FecR) were further transformed with pSTD689 (vec), pYH9 (RseP-HM, WT) or pYH13 (RseP(E23Q)-HM, E23Q). The cells were grown as in Fig. 3C. The proteins were analyzed by 10% Laemmli SDS-PAGE and anti-FLAG immunoblotting or 12.5% Laemmli SDS-PAGE and anti-RseP (α-RseP), or anti-SecB (α-SecB) immunoblotting. The band intensity of RseP-HM is saturated. (C) Behaviors of the truncated derivatives of F-MBP-FecR. YK191 cells harboring pSTD689 (vec), pYH9 (WT), or pYH13 (E23Q) were further transformed with pYK147 derivatives encoding the truncated mutant forms of F-MBP-FecR shown in (A). The cells were grown and analyzed as in (B). The lanes with an asterisk were the same sample used for the comparison of the band sizes. Note that the intensities of RseP-HM and some of F-MBP-FecR and its derivative CL(a), CL(b) and CL(c) bands are saturated.

To estimate the approximate position of the C-terminal end of the CL(a), CL(b), and CL(c) fragments, we constructed a series of C-terminally truncated versions of F-MBP-FecR (Fig. 4A). The transmembrane region of FecR was predicted to be located between residues 80 and 100 (with the N-terminal Met residue of FecR set to residue 1) by the topology prediction algorithm SPOCTOPUS (http://octopus.cbr.su.se/index.php)(61). Accordingly, we constructed two truncated mutants, F-MBP-FecR101 and F-MBP-FecR79, in which the regions after residues 101 and 79 had been deleted, respectively. F-MBP-FecR79 lacks the predicted transmembrane and periplasmic regions, whereas F-MBP-FecR101 only lacks the periplasmic region. Because an alternative region (residues 86 to 100) of FecR has been suggested to span the membrane in a previous study (42), we also constructed a truncated mutant (F-MBP-FecR85) that lacked the region after residue 85. In addition, based on the previous studies that reported the self-cleavage of FecR between Gly-181 and Thr-182 (59, 62, 63), we constructed F-MBP-FecR181 without the region after residue 181, as a mimic of the N-terminal fragment generated by self-cleavage. Finally, we systematically constructed three additional C-terminally truncated mutants (F-MBP-FecR161, 141, and 121) by deleting C-terminal 20, 40, or 60 residues from F-MBP-FecR181, respectively. These constructs were expressed in Δ*rseA* Δ*rseP* cells and their sizes were compared with the CL(a), CL(b) and CL(c) fragments that had been generated from full-length F-MBP-FecR *in vivo* (Fig. 4C). With F-MBP-FecR181, two bands having nearly the same mobilities as CL(a) and CL(b), respectively, were detected (Fig. 4C, compare *lanes 1-3* with *4*). The former band would be intact in F-MBP-FecR181, suggesting that CL(a) is a self-cleavage product of F-MBP-FecR (59, 62, 63). Similarly, two bands were observed with F-MBP-FecR161, 141, and 121 (Fig. 4C, *lanes 4-7*), and the upper bands would also represent their “intact” form. The second (smaller) bands generated for the above four constructs had almost the same apparent size as CL(b). In contrast, only a single band was detected for F-MBP-FecR101, 85, and 79 (Fig. 4C, *lanes 8-10*), each having no periplasmic region. Taken together, these results strongly suggest that FecR receives a cleavage in its periplasmic region to generate CL(b). As the “intact” band of F-MBP-FecR121 was slightly larger than the CL(b) band, the possible cleavage site would be located just upstream of reside 121.

The mobility of CL(c), whose production is dependent on the proteolytic function of RseP, was almost the same as that of F-MBP-FecR85 (compare Fig. 4C, *lanes 8-10* and *11*), suggesting that CL(c) would be generated by the RseP-mediated cleavage around residue 85. This cleavage site is highly likely to be located within the transmembrane region of FecR, given that RseP cleaves a transmembrane segment of a substrate. Accordingly, our results support the prediction that the FecR spans the membrane in the region between residues 80 and 100.

### RseP cleaves the FecR CL(b) fragment and converts it to the CL(c) fragment

The apparent RseP-dependent conversion of CL(b) to CL(c) can be well-explained if CL(c) is produced by the intramembrane proteolysis of CL(b) by RseP. To substantiate this possibility, the stability and the production kinetics of F-MBP-FecR and its derivatives (CL(a), CL(b) and CL(c)) were investigated by pulse-chase experiments (Fig. 5 and S3). F-MBP-FecR was expressed in the Δ*rseP* strain, labeled with [^35^S]methionine and chased with unlabeled methionine up to 81 min after initiation of the chase. FL and CL(a) band were the only species detected just after the start of the chase (Fig. 5A, and the close-up view of the quantified results for the early periods of the pulse-chase experiments is shown in Fig. S3B, at 0.25-min time point). FL almost completely disappeared within 3-min chase. On the other hand, CL(a) decreased gradually with time with a small amount detectable even after the 81 min-chase (Fig. 5B). In contrast, CL(b) appeared at 1-min chase point and increased during the chase period. Co-expression of wild-type RseP-HM only marginally accelerated the disappearance of FL and CL(a). In this case, however, CL(b) was not detected, but instead CL(c) was generated. CL(c) increased with a similar kinetics to that of CL(b) observed in the absence of the RseP-HM co-expression. Generation of CL(c) was dependent on the proteolytic function of RseP-HM as co-expression of RseP(E23Q)-HM did not exert such an effect, although the accumulation levels of RseP-HM and RseP(E23Q)-HM were comparable (Fig. S3A). These results strongly suggest that both CL(b) and CL(c) are derived from CL(a). However, it was still unclear from these results whether CL(c) was generated from CL(a) or generated from CL(b).

**Figure 5.**
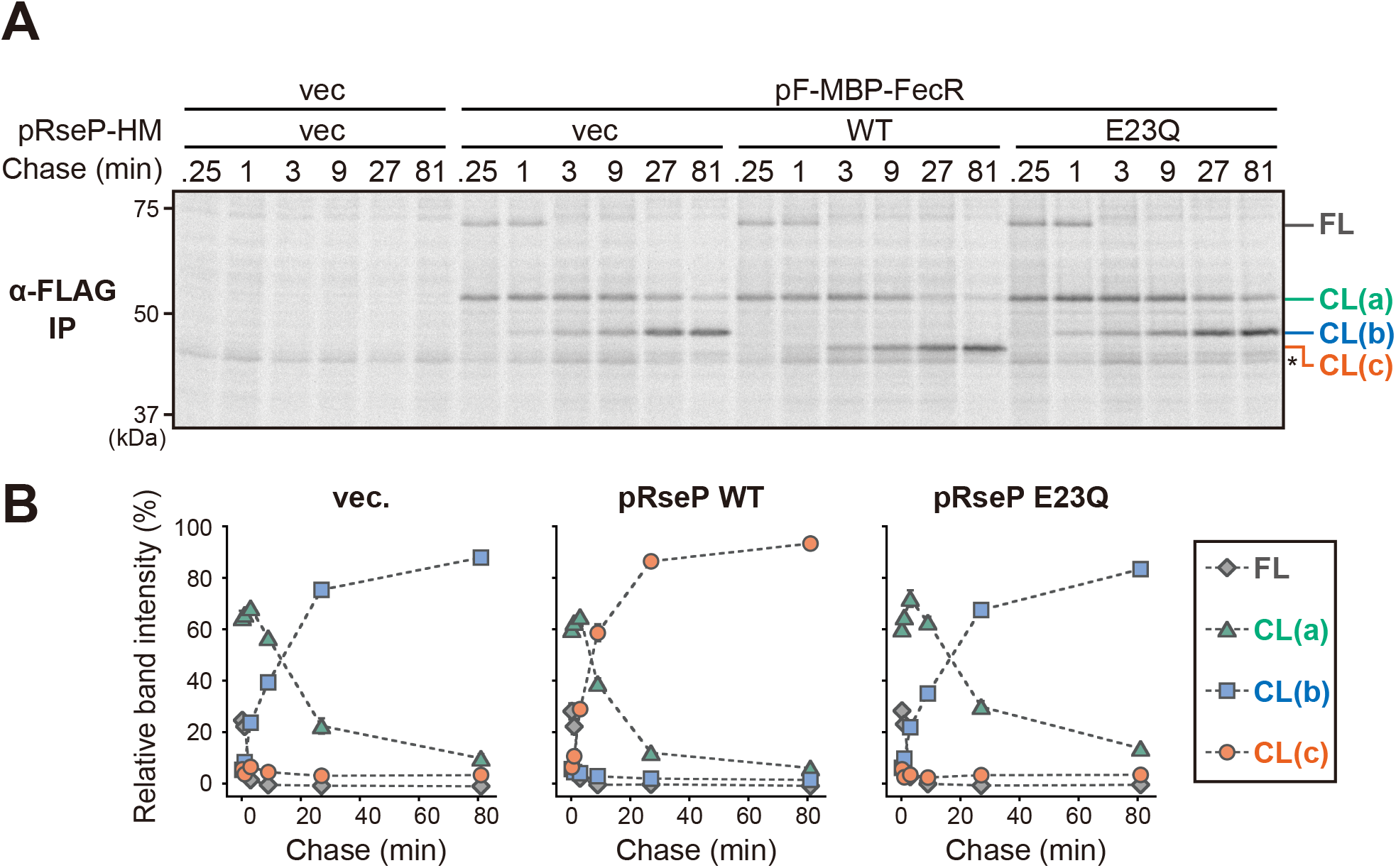
Kinetics and RseP-dependence of the generation of the FecR-derived fragments. Pulse-chase analysis of the F-MBP-FecR cleavage by RseP. YK191 (Δ*rseA* Δ*rseP*) cells harboring pTWV228 (vec) or pYK147 (pF-MBP-FecR) were further transformed with pSTD689 (vec), pYH9 (RseP-HM, WT), or pYH13 (RseP(E23Q)-HM, E23Q). The cells were grown at 30°C in M9-based medium containing 10 μM FeCl3, 1 mM Na3-citrate until early-log phase. After induction with 1 mM IPTG and 1 mM cAMP for 10 min, proteins were pulse-labeled with [^35^S]methionine for 1 min followed by chase with an excess amount of unlabeled methionine for the indicated periods. The proteins were analyzed by anti-FLAG immunoprecipitation, 10% Laemmli SDS-PAGE, and phosphorimaging. Asterisk indicates a non-specific background band. Quantification of the experimental results shown in (A). The percentage of the intensity of each F-MBP-FecR fragment in total band intensities was plotted versus the chase time. The averages of the values from two independent experiments are shown with standard deviations. Because the exact number of the Met residue in the cleaved fragments is not known, the band intensities were not corrected by the number of Met.

To address this point, we designed an assay procedure (Figs. 6A,6B and S4) in which cells expressing F-MBP-FecR from a constitutive expression vector were pulsed-labeled and chased for 5.5 min, and then, RseP-HM or RseP(E23Q)-HM was induced (Fig. 6A, B). At 0.5 min before the induction of RseP, only CL(a) and CL(b) were detected. We expected that, if RseP is indeed involved in the cleavage of CL(b), (i) this experimental system could enable us to detect the possible RseP-dependent conversion of pre-existing CL(b) to CL(c), and also (ii) the presence of an excess amount of the unlabeled FL and CL(c) molecules (as the result of their constitutive expression) compared with RseP-HM that could retard apparent degradation rate of labeled CL(b) and facilitate detection of the conversion process. Actually, we captured the CL(b) to CL(c) conversion upon the induction of RseP-HM but not that of RseP(E23Q)-HM. These results strongly support the idea that RseP directly cleaves CL(b) to produce CL(c). Although the accumulation levels of RseP-HM and RseP(E23Q)-HM varied significantly in the two independent experiments (Fig. S4B), the results of the pulse-chase analysis of the F-MBP-FecR derivatives were highly reproducible (Fig. 6B and Fig. S4A). Although the reason of this observation remains unclear, it might suggest that only a limited portion of overexpressed RseP acts in the CL(b) cleavage.

**Figure 6.**
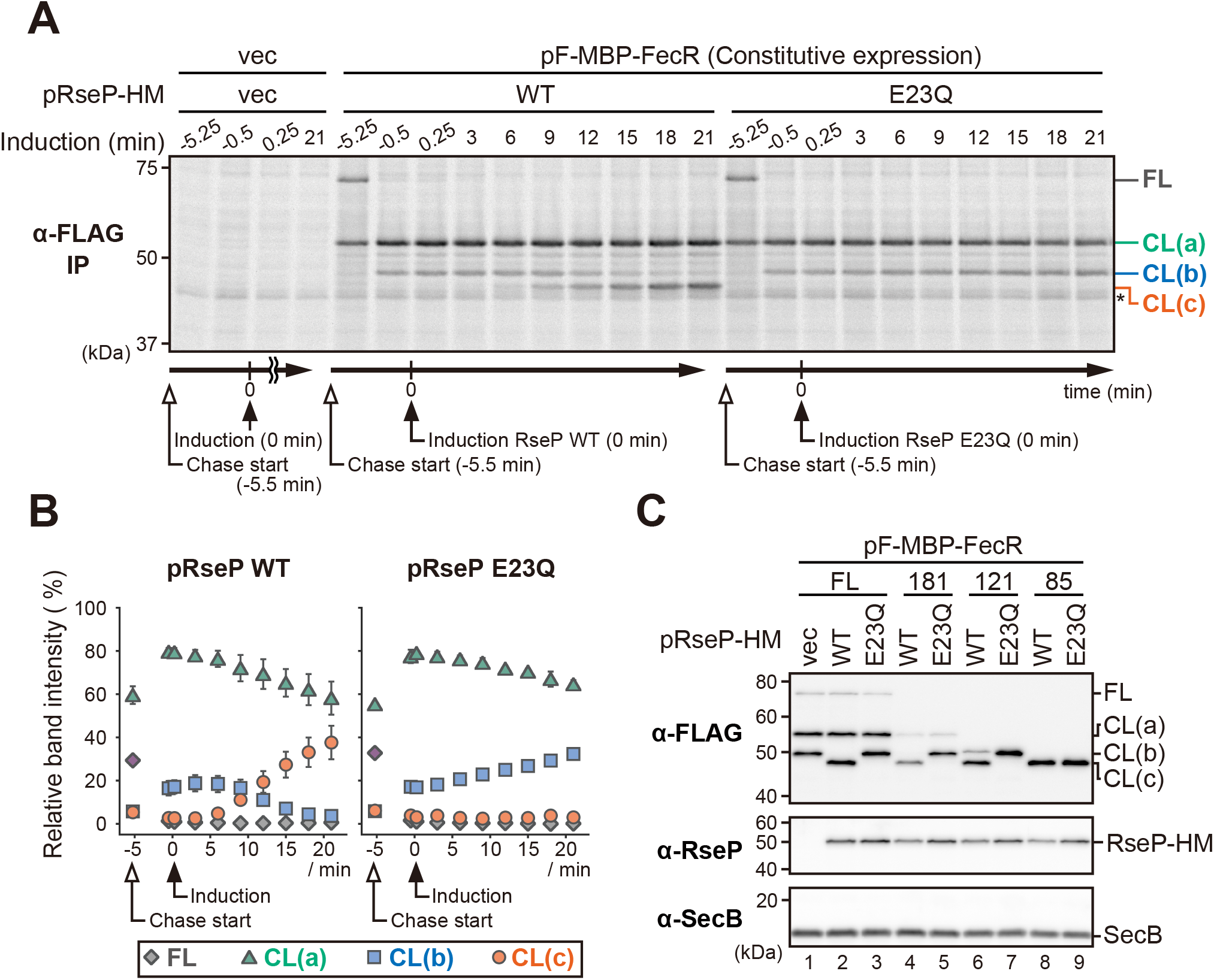
RseP-dependent conversion of the FecR CL(b) fragment to CL(c). (A) Pulse-chase analysis of the RseP-dependent cleavage of F-MBP-FecR. YK191 (Δ*rseA* Δ*rseP*) cells harboring pYH604 (constitutive expression vector pTWV228c, vec) or pYK182 (pF-MBP-FecR) were further transformed with pSTD689 (vec), pYH9 (RseP-HM, WT) or pYH13 (RseP E23Q-HM, E23Q). The cells were grown and analyzed as in Fig. 5A except that F-MBP-FecR was expressed constitutively and that RseP was induced after the start of the chase. Following pulse-labelling with [^35^S]methionine for 1 min, chase was initiated by the addition of unlabeled methionine. The RseP proteins were induced with 1.2 mM IPTG and 1.2 mM cAMP 5.5 min after the initiation of the case (indicated as 0 min). The timing of the chase-start and the RseP induction was shown under the gel image. (B) Quantification of the results of the experiments shown in (A). The percentage of each band intensity to the total band intensity are plotted. (C) Behaviors of the truncated derivatives of F-MBP-FecR in response to the proteolytic activity of RseP. YK191 cells harboring pSTD689 (vec), pYH9 (WT) or pYH13 (E23Q) were further transformed with a plasmid encoding the F-MBP-FecR derivatives as in Fig. 4A. The cells were grown and analyzed as in Fig. 4B.

To confirm the direct involvement of RseP in cleavage of CL(b), we examined the effect of RseP on the degradation of the C-terminally truncated mutants of F-MBP-FecR. F-MBP-FecR121, which lacked almost the entire periplasmic region, had a slightly larger size than that of CL(b) and generated CL(b) but not CL(a). The model proteins were co-expressed with wild-type or the E23Q mutant form of RseP-HM (Figs. 4A and 6C). CL(b) generated from F-MBP-FecR121 and F-MBP-FecR181 were converted to CL(c) in the functional RseP-dependent manner. (Fig. 6C, *lanes 4*-*7*) On the other hand, the band pattern and accumulation of F-MBP-FecR85, which mimics the CL(c) fragment, was little affected by the RseP function (Fig. 6C, *lanes 8* and *9*). These results further support the RseP-catalyzed cleavage of CL(b) to yield CL(c).

### The FecR CL(b) fragment is generated from the CL(a) fragment

CL(b) was generated from F-MBP-FecR181 that mimics CL(a) (Fig. 4C *lane 4* and Fig. 6C *lane 5*). It would be thus conceivable that CL(b) is a degradation product of CL(a). Our pulse-chase results, showing the apparent precursor-product relationship between CL(a) and CL(b) (that is, the disappearance of CL(a) is accompanied by the appearance of CL(b)) (Fig. 5), also strongly support that CL(a) is converted to CL(b). Immunoblotting analysis showed that the addition of citrate to the medium led to the increase in the accumulation of CL(b) and the concomitant decrease in that of CL(a) (Compare Fig. 3C *lanes 6* with *10* (F-FecR), or Fig. 4B *lanes 6* with *10* (F-MBP-FecR; the quantified results of Fig. 4B are shown in Fig. S2). These results suggest that the CL(a) to CL(b) conversion is promoted by the ferric citrate signal.

### The essential function of RseP in the transcriptional activation of the *fec* operon is to produce the FecR CL(c) fragment

The above results showed that RseP proteolytically converts FecR CL(b) to CL(c). The generation of CL(c) was well-correlated with the ferric citrate-dependent activation of the *fec* operon (Fig. 2 and 3). In addition, a previous study showed that the expression of the predicted cytoplasmic domain of FecR (FecR(1-85)) (40, 42, 64), which mimics CL(c), activated the transcription of the *fec* operon with no need of the ferric citrate signal. These results strongly suggest that RseP activates the *fec* operon by producing the CL(c) fragment. However, it still cannot be excluded that RseP plays an additional role in the activation of the *fec* operon. We thus investigated the requirement of RseP in the *fec* operon activation by the cytoplasmic fragment of FecR (Fig.7). When expressed in the Δ*fecR* strain, F-FecR85, a 3xFLAG-tagged cytoplasmic domain (residue 1 to 85) of FecR showed almost the same mobility as that of CL(c) generated from F-FecR (Fig. 7A, compare *lanes 5* and *6* with *4*), further supporting that CL(c) is generated by the cleavage around the residue 85. F-FecR85 accumulated at similar levels in the presence or absence of citrate (Fig. 7A, *lanes 5* and *6*). Also, the expression of F-FecR85, in contrast to that of full-length F-FecR, activated the *fec* operon transcription in a citrate-independent manner (Fig. 7B, pF-FecR). The N-terminally attached 3xFLAG tag little affected the function of FecR and FecR85 as the un-tagged versions of these proteins gave essentially the same results (Fig. 7B, compare pFecR with pF-FecR). LacZ activities showed apparent correlations with the accumulation levels of F-FecR85 or F-FecR CL(c) fragment (Fig. 7A, *lanes 3-6* and Fig. 7B, pF-FecR). These results confirmed the previous finding that the cytoplasmic domain of FecR can constitutively activate the transcription of the *fec* operon (40, 42, 64). The RseP function was not required for this ferric citrate-independent activation of the *fec* operon (Fig. 7C), as the reporter activities of the cells expressing F-FecR85 were elevated considerably in a citrate-independent manner both in the presence or absence of chromosomal *rseP*. Consistent with the above result, the accumulation level of F-FecR85 in the *rseP*^+^ and Δ*rseP* cells was not affected by the addition of citrate (Fig. S5). In this experiment, the reporter activities in the Δ*rseP* cells were apparently a little higher than those in the *rseP*^+^ cells, possibly reflecting the increased accumulation of the F-FecR85 protein in the former cells (Fig. S5), although the exact reason for this differential accumulation is unclear. Taken together, these results demonstrated that the essential role of RseP in the activation of the *fec* operon is to produce the CL(c) fragment that co-functions with FecI from CL(b).

**Figure 7.**
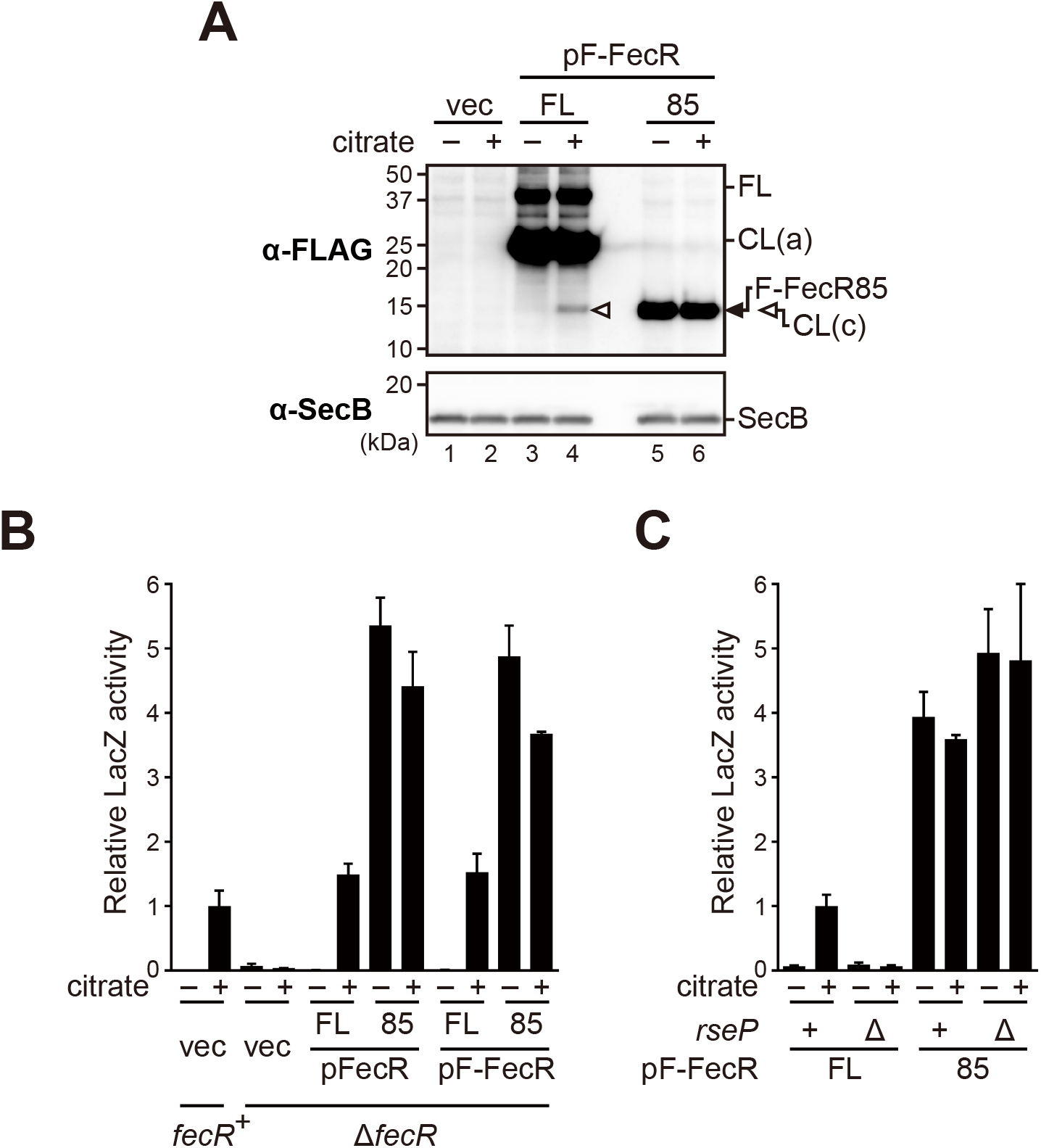
The CL(c) fragment can activate the expression of the *fec* operon in an RseP-independent manner. (A) The behaviors of F-FecR and F-FecR85 in response to the addition of Na3-citrate. MC4100 (*fecR*^+^) or YK627 (Δ*fecR*) cells harboring pYK149 (PfecA-*lacZ*) and pSTD343 (*lacI*) were further transformed with pSTD1060 (vec), pYK186 (pFecR, FL), pYK198 (pFecR85), pYK188 (pF-FecR, FL), or pYK200 (pF-FecR85). The cells were grown as in Fig. 2B and proteins were analyzed as in Fig. 3C. Note that the band intensities of F-FecR FL and CL(a), and F-FecR85 are saturated. (B) The citrate-dependency of the *fec* operon activations mediated by the FecR and F-FecR derivatives. The cells are grown and the LacZ activity was measured as in Fig. 3B. (C) The RseP-dependency of the *fec* operon activations mediated by F-FecR85. YK630 (*rseP*^+^) or YK797 (Δ*rseP*) cells harboring pYK149 (PfecA-*lacZ*) and pSTD343 (*lacI*) were further transformed with pYK188 (pF-FecR, FL) or pYK200 (pF-FecR85). The cells were grown and the LacZ activity was measured as in (B). The relative LacZ activities were normalized by the LacZ activity of the cells expressing F-FecR that had been grown in medium containing 1 mM Na3-citrate are shown.

## Discussion

In this study, we performed proteomic analysis to identify the substrates of RseP, the S2P family intramembrane protease of *E. coli*, to explore novel functions of this protease, and unveiled that RseP is involved in the regulation of the Fec system (Ferric citrate uptake system) genes through the intramembrane cleavage of a novel physiological substrate, FecR. Our proteomic analysis identified multiple Fec system components (FecA, FecD and FecE) encoded by the *fec* operon as proteins whose levels were significantly decreased in the RseP-deficient strain, suggesting that RseP is required for the expression of the *fec* operon. We thus examined the transcriptional regulation of this operon by using a *lacZ*-reporter and demonstrated that proteolytic activity of RseP is essential for its activation. Transcription of this operon is known to be controlled by FecI, an alternative sigma factor (36–39), whose activity is regulated by a cytoplasmic membrane protein, FecR (40–42). Since FecR is a single-pass transmembrane protein of type II topology, which is a shared feature of substrates of the bacterial S2P proteases including RseP, we examined the possibility that RseP cleaves this protein to induce expression of the *fec* operon. The experiments using the FecR-derived model substrates revealed that FecR receives sequential processing at the membrane and that RseP participates in the last step in the processing that generates the cytoplasmic tail fragment of FecR required for the transcriptional activation of the *fec* operon.

FecR was processed to yield the fragments that we named here CL(a), CL(b), and CL(c) (Fig. 5). We demonstrated that RseP cleaves the transmembrane region of CL(b), and converts it to CL(c) (Fig. 6). The production of CL(c) is required and sufficient for the FecI-mediated activation of the *fec* operon transcription (Fig. 2 and 7). These observations coincide with and further support the results of previous studies, which showed that the expression of the cytoplasmic region of FecR led to the ferric citrate-independent activation of the transcription of the *fec* operon (40, 42, 64). A previous study showed that the N-terminal cytoplasmic region of FecR can directly interact with FecI (60) and acts as a “pro-sigma” factor to facilitate the transcription of the *fec* operon. It is thus suggested that the RseP-mediated cleavage of CL(b) would liberate CL(c) from the membrane to activate the *fec* operon transcription by forming a complex with FecI. Braun *et al*. (2006) previously proposed a model wherein the RseP-catalyzed intramembrane proteolysis of FecR is involved in the activation of the *fec* operon (65), although it has not been demonstrated experimentally. Our present study provided the strong evidence supporting this model. RseP orthologues in gram-negative bacteria act in processing of sigma factor regulators to activate cognate sigma factors in a variety of iron acquiring systems (*e.g.*, *B. bronchiseptica* HurR (21), *P. aeruginosa* FpvR (20), FoxR (63), HxuR, HasS (66) and *P. putida* IutY (51)). Our study added another example to this list.

Our results suggest that CL(b) is produced by the cleavage of CL(a) and that this process is promoted by the ferric citrate signal. Thus, CL(a) apparently receives two successive cleavages to yield CL(c), which is reminiscent of the two-step cleavage of *E. coli* RseA triggered by the extracytoplasmic stresses. In case of RseA, it is recognized and cleaved by RseP (“site-2 cleavage”) only after it received prior “site-1 cleavage” by the membrane serine protease DegS on the periplasmic side. While the two successive cleavages are common among many bacterial S2P substrates, the protease catalyzing the site-1 cleavage is not necessarily a DegS homolog. For example, *B. subtilis* PrsW, a multipass metalloprotease that cleaves RsiW in response to antimicrobial peptides and envelope stresses is unrelated to *E. coli* DegS (67). Also, in case of *P. putida* IutY, periplasmic serine protease Prc is possibly responsible for its “site-1 cleavage” (51). Although we have no information on the putative protease(s) responsible for the production of CL(b) form CL(a) in the FecR-mediated signal transduction, the ferric citrate signal might induce some conformational change in CL(a) to make it susceptible to the site-1 cleavage. Alternatively, the ferric citrate signal might directly activate the putative “site-1” protease. Further analysis, especially the identification of the putative “site-1” protease, will be needed to reveal the detailed molecular mechanism of this process.

The production of CL(a) occurred immediately after the synthesis of FecR and independently of RseP. A previous study suggested that CL(a) is generated by the cleavage between Gly-181 and Thr-182 (59), which is consistent with our result that CL(a) migrated on an SDS-PAGE gel to almost the same position as the FecR-derivative that had been truncated at Gly-181. This GT motif is conserved among several anti-sigma factors of *P. aeruginosa* and *P. putida* (63) and *E. coli* FecR. Furthermore, Bastiaansen *et al.* (2015) showed that one of them, *P. aeruginosa* FoxR, underwent self-cleavage at this site and suggested that the other GT motif containing anti-sigma proteins are also processed auto-catalytically at this motif in the non-enzymatic fashion, *i.e.*, N-O acyl rearrangement (62). It would be very likely that *E. coli* FecR CL(a) is generated by the self-cleavage at the GT motif, too. Whether prior autoproteolysis is required for the production of CL(b) is unclear. Previous studies showed that some mutations in the GT motif of FoxR, which receives a cleavage by RseP during the signal transduction like FecR, blocks the self-cleavage but still allow iron-dependent signal transduction, suggesting that the self-cleavage is not essential in this system, although it cannot be excluded that the FoxR mutants with the GT motif mutations receive degradation by some proteases to generate a small amount of a CL(a)-like fragment, which is sufficient to drive the down-stream reactions. If the self-cleavage of FecR is not required for the signal transduction in the *E. coli* Fec system as suggested for FoxR, the possible protease responsible for generation of CL(b) might be able to cleave both of the FL and CL(a) forms of FecR to provide a substrate of RseP, as RseP should not cleave full-length FecR like RseA discussed below.

We propose a model of the FecR-processing cascade in the signal transduction by the Fec system (Fig. 8) based on the results of preceding studies by other researchers and the present study. (i) FecR is self-cleaved almost immediately after synthesis and generates CL(a). (ii) In response to the ferric citrate signal transmitted via FecA, CL(a) receives the C-terminal truncation or trimming by unknown protease(s) in the periplasmic space to generate CL(b). (iii) CL(b) is then recognized and cleaved by RseP within the membrane, leading to the release of CL(c) from the membrane. (iv) Finally, the liberated CL(c) that is associated with the FecI promotes the transcription of the *fec* operon. The approaches used in this study, especially detailed pulse-chase experiments and analyses with various model proteins, will be useful to understand the details of the processing of other proteins including *P. aeruginosa* FoxR for which similar multi-step cleavages have been proposed but their timings and relationships are not fully understood. Regulated intramembrane proteolysis (RIP) catalyzed by intramembrane proteases is widely found in many organisms from bacteria to eukaryote (1, 2). The cleavage of FecR to induce the *fec* operon transcription in the signal-dependent activation of the Fec system would be one of the typical instances of RIP-mediated transmembrane signaling.

**Figure 8.**
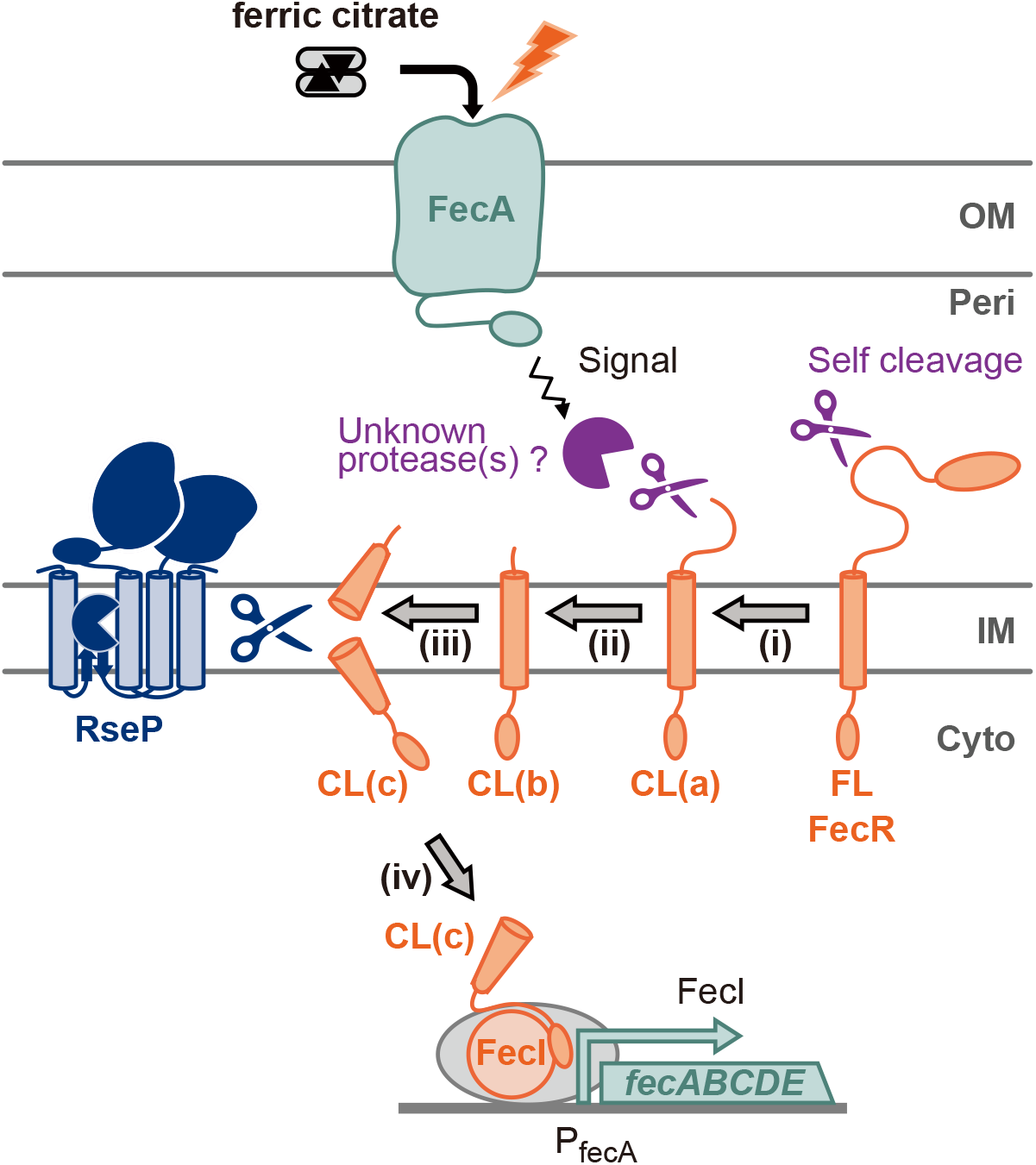
A model of the sequential processing of FecR and the involvement of RseP in it in the ferric citrate-induced signal transduction. (i) FecR undergoes a self-cleavage immediately after translation to generate CL(a). (ii) CL(a) is cleaved or trimmed by unknown protease(s) in the periplasm to yield CL(b), which is promoted by the ferric citrate signal. (iii) CL(b) then receives RseP-mediate intramembrane proteolysis inside the membrane to release CL(c) from the membrane. (iv) sigma factor FecI bound to CL(c) activates the transcription of the *fec* operon.

We previously proposed that the periplasmic PDZ tandem of RseP serves as a size-exclusion filter to avoid proteolysis of membrane proteins with a large periplasmic domain by RseP (27). In the σ^E^ extracytoplasmic stress response, the site-1 protease DegS is activated by the stress signals and cleaves full-length RseA, the ant-σ^E^ protein, in its periplasmic region, which triggers the following site-2 cleavage by RseP as the RseA degradation intermediate (RseA148) generated by the site-1 cleavage has a small periplasmic region (~30 a. a.) and can get access to the intramembrane active site passing through the PDZ filter. Thus, the PDZ filter ensures the stress dependent activation of σ^E^ (Fig. S6) (68, 69). Similarly, in the case of FecR we expect that the conversion of CL(b), which is supposed to have a periplasmic region of about 80 a. a., to CL(c) with a smaller periplasmic tail (~ 20 a. a.) would make FecR accessible to the intramembrane active site of RseP through the PDZ filter. As the CL(b)-to-CL(c) conversion is promoted by the ferric citrate, the PDZ filter-mediated discrimination of the substrates contributes to the ferric citrate signal-dependent activation of FecI as well.

The predicted transmembrane sequence of FecR (R^80^HVMKGLLLLLGAGGGWQLWQ^100^) has no apparent sequence homology to the known substrates of RseP (14, 28, 29). We previously suggested that the stability of a transmembrane helix in the membrane, but not the exact primary sequence, is an important determinant for its susceptibility to cleavage by RseP (28, 29). Consistent with this notion, RseA and FecR have several residues such as Lys, Arg, and Gln that potentially destabilize a helix in a hydrophobic environment (70) in their predicted transmembrane segments. These residues might be required for the efficient cleavage of these proteins by RseP. In addition, similar to RseA, the transmembrane sequence of FecR has relatively low hydrophobicity compared with transmembrane segments of typical membrane proteins such as lactose permease (LacY). The assigned transmembrane sequences of FecR and RseA are not predicted as a transmembrane segments by a transmembrane prediction program such as TMHMM (71) in contrast to the transmembrane segments of LacY (Fig. S7). The products of intramembrane proteolysis could be released from the membrane because they have a shorter hydrophobic stretch and the polarity or charges at their N- and C-termini generated by the intramembrane cleavage (72). The low hydrophobicity of the transmembrane segments of RseA and FecR could further facilitate efficient liberation of the N-terminal fragments of RseA and FecR from the membrane after the RseP-mediated intramembrane proteolysis and accelerate the transmembrane signaling in each system. Interestingly, the first and fifth transmembrane segments of LacY can be efficiently cleaved by RseP when placed in a model substrate (29) despite of their high hydrophobicity, suggesting that the overall hydrophobicity of a transmembrane segment is not a critical determinant for its susceptibility to RseP.

Proteomic analysis is a powerful approach that allows non-biased search for protease substrates. Thus far, many examples have been reported in which proteomic analysis was successfully used to identify the substrate of membrane proteases including *E. coli* FtsH (32), *B. subtilis* MgtE (30), *Homo sapiens* PARL (33), etc. We thus employed this approach and identified FecR as novel substrate of RseP. In this analysis we found significant changes in the accumulation levels of some proteins, in addition to the Fec proteins, depending on the proteolytic activity of RseP. Among the group of proteins that exhibited increased accumulation in the RseP-deficient strain, CyoE (73), Fiu (74), CirA (75), and FepA (76) have been reported to be negatively regulated by the Fur protein. Fur acts as a global repressor for the expression of genes involved in iron uptake upon its interaction with iron (77). The increased expression of the Fur-regulated genes in the cells expressing RseP(E23Q) might at least partially be ascribed to a decreased intracellular iron level resulting from the lowered expression of the Fec components. In contrast to the above-mentioned proteins, the increased accumulation was observed for AmtB and ZupT (fold change >2) in the cells expressing RseP(E23Q) (Fig. 1 and Table. 1). Both AmtB and ZupT are transporters of the cytoplasmic membrane; the former facilitates the uptake of ammonia (48, 78) whereas the latter imports divalent metal cations (49, 50). Since these proteins are multi-spanning transmembrane proteins, it is unlikely that RseP directly proteolyze these proteins. Their expression or stability might be indirectly affected by the loss of the proteolytic function of RseP. It should be noted that FecR evaded detection in our proteomic analyses. One possibility would be that the FecR level was below the detection threshold in our experiments. Another possibility would be that most of the FecR molecules were lost during the preparation of the membrane fraction. As discussed above, FecR and RseA have a transmembrane segment of relatively low hydrophobicity. We often encountered the significant decrease or loss of RseA and its derivatives in the membrane fraction possibly as a result of their falling-out from the membrane (unpublished observations). It is possible that we overlooked other RseP substrates by similar and/or other reasons. For the systematic identification of RseP substrates, it is necessary to improve the methods of sample preparations in mass analyses (31, 79, 80) and also use additional approaches such as the co-isolation of possible substrates with a proteolytically inactive RseP and systematic screening of type II membrane proteins in combination with the mass analysis-based proteomic approach. Such approaches would also be useful to search for the unknown substrate of other IMPs.

Recently, the Fec system has been reported to be crucial in bovine mastitis caused by a pathogenic *E. coli* strain (81). Bovine mastitis is a disease in dairy cows that causes economic loss to the global dairy industry. This study and further analysis of the regulation of the Fec system may hopefully contribute to unveiling the mechanism and development of the treatment of this disease. Our mass analysis results suggest that the levels of the proteins involved in a variety of cellular activities such as ion transport, rRNA processing, and acetylation are affected by the impairment of the RseP function (Table S1). It raises the possibility that RseP may have additional substrates and act directly or indirectly in still unknown cellular processes. Further study by using a variety of approaches discussed above will lead to the comprehensive understanding of the significance of RseP in cellular activities.

## Experimental procedures

### Media

L broth (10 g/L Bacto Tryptone, 5 g/L yeast extract and 5 g/L NaCl; pH adjusted to 7.2 by using NaOH) and M9 medium (without CaCl_2_) (82) supplemented with 2 μg/mL thiamine and 0.4% glucose were used for cultivation of *E. coli* cells. Ampicillin (50 μg/mL), chloramphenicol (20 μg/mL) and/or spectinomycin (50 μg/mL) were added for selecting transformants and for growing plasmid-harboring cells. Bacterial growth was monitored using mini photo 518R (660 nm; TAITEC) or Klett-Summerson colorimeter (filter no. 54; Klett Manufacturing).

### Strains, plasmids and oligonucleotides

*Escherichia coli* K-12 strains, plasmids and oligonucleotides used in this work are listed in Supplementary Tables S2, S3 and S4, respectively. Construction of the individual strains and plasmids are described in Supplementary Experimental Procedures.

### Antibodies

Monoclonal anti-FLAG M2 antibody (MilliporeSigma), Rabbit polyclonal anti-RseP antibody (83) and anti-SecB antibody (26) were used for immunoblotting. Monoclonal anti-FLAG M2 Affinity Gel (MilliporeSigma) was used for Immunoprecipitation in pulse-chase assay.

### NanoLC/MS/MS sample preparation

Cells were grown in L broth supplemented with 0.4% glucose, 1 mM IPTG and 1 mM cAMP at 30°C until a mid-log phase. Harvested cells were washed with 10 mM Tris-HCl (pH 8.1) and suspended in buffer containing 10 mM Tris-HCl (pH 8.1) and protease Inhibitor cocktail (Nacalai Tesque). Cell suspension was frozen, thawed, and lysed by sonication. After cell debris was removed by low speed centrifugation, membrane fractions were collected by ultracentrifugation (at 125,000 × *g*, 60 min) and suspended in PTS solution (12 mM Sodium deoxycholate, 12 mM sodium N-lauroylsarcosinate, 100 mM Tris-HCl (pH 9.0)) (84). After incubation at room temperature for 30 min and at 37°C for 5 min, the protein amount was confirmed with a BCA protein assay kit (Thermo Fisher Scientific). Proteins were reduced by treatment with 10 mM DTT for 30 min, alkylated by treatment with 50 mM chloroacetamide for 30 min in the dark. Next, the protein solution was 5-fold diluted with 50 mM ammonium bicarbonate and the proteins were digested with Lys-C (w/w 1:100) for 3 h, followed by trypsin digestion (w/w 1:100) overnight at 37 °C. The peptides were desalted using StageTip (85) with SDB-XC Empore disk membranes (GL Sciences). The peptides were labeled with Tandem Mass Tag (TMT) (Thermo Fisher Scientific), fractionated using strong cation exchange chromatography (SCX) StageTips (86) and suspended in the loading solution (0.5% TFA and 4% ACN) for subsequent nanoLC/MS/MS analyses.

### NanoLC/MS/MS analyses

NanoLC/MS/MS analyses were performed on an Orbitrap Fusion Lumos (Thermo Fisher Scientific), connected to an Ultimate 3000 pump (Thermo Fisher Scientific) and an HTC-PAL autosampler (CTC Analytics). Peptides were separated on a self-pulled needle column (150 mm length × 100 μm ID, 6 μm opening) packed with Reprosil-C18 AQ 3 μm reversed-phase material (Dr. Maisch). The flow rate was set to 500 nL/min. The mobile phase consisted of (A) 0.5% acetic acid and (B) 0.5% acetic acid in 80% acetonitrile. Three-step linear gradients of 5-10% B in 5 min, 10-40% B in 60 min, and 40-100% B in 5 min were employed. For TMT-labeled samples, synchronous precursor selection-MS3 (SPS-MS3) (87) was performed. The MS scan range was *m/z* 375-1500. MS scans were performed by the Orbitrap with *r* =120,000, MS/MS scans were performed by the Ion Trap in Turbo mode and MS3 scans were performed by the Orbitrap with *r* = 15,000. Auto gain control was set to 4.00 × 10^5^, 1.00 × 10^4^, 5.00 × 10^4^ for MS, MS/MS and MS3, respectively. The normalized CID collision energy was set to 35.

### Database searching

The raw MS data files were analyzed by MaxQuant v1.6.2.3 (88). Peptides and proteins were identified by means of automated database searching using Andromeda against the *E. coli* K-12 SwissProt Database (version 2018-07, 4,324 protein entries) with a precursor mass tolerance of 20 ppm for first search and 4.5 ppm for main search and a fragment ion mass tolerance of 0.5 Da Enzyme was set as trypsin/P (cleaves after lysine and arginine also if a proline follows) and semi-specific search was performed. Cysteine carbamidomethylation was set as a fixed modification. Methionine oxidation and acetylation on protein N-termini were set as variable modifications. The search results were filtered with FDR < 1% at the peptide spectrum match (PSM) and protein levels.

### *β*-galactosidase (LacZ) activity assay

LacZ activity of the cells carrying the reporter plasmid pYK149 (PfecA-*lacZ*) was measured basically according to the procedure described previously (89). The cells were grown at 30°C in M9-based medium with 20 μg/mL each of the 20 amino acids, 2 mg/mL thiamine and 0.4% glucose until mid-log phase. The cells were mixed with Reporter 5xLysis buffer (Promega), frozen at −80°C for more than 1 h, and thawed by incubation at 37°C for 30 min in a clear 96-well plate. Then, the equal volume of Z-buffer (60 mM Na_2_HPO_4_·7H_2_O, 40 mM NaH_2_PO_4_·H_2_O, 10 mM KCl, 1 mM MgSO_4_·7H_2_O, 40 mM *β*-mercaptoethanol) containing 1.32 mg/mL 2-nitrophenyl *β*-D-galactopyranoside (ONPG, MilliporeSigma) was added to this lysate and incubated at room temperature, followed by detection of the absorbance at 420 and 550 nm at every 2 min using the Viento Nano microplate reader (BioTek Instruments). The relative LacZ activity was calculated as follows. First, the "raw LacZ activity" was calculated according to the following equation; "raw LacZ activity (arbitrary units)" = (A420 - 1.75 × A_550_) / (incubation time (min)). The value of the raw LacZ activity of each sample was divided by that of a standard cell sample (CU141 cells cultured in M9-based medium), and then by A600 of the bacterial culture at the time of collection, giving the "corrected LacZ activity". Finally, the relative LacZ activity of each sample was obtained by dividing the value of the corrected LacZ activity by that of the corresponding control (see the legends for the control in each experiment).

### SDS-PAGE

Proteins were dissolved in SDS sample buffer (62.5 mM Tris-HCl (pH 8.5), 2% (w/v) SDS, 10% glycerol, 10% *β*-mercaptoethanol, trace of bromophenol blue) and separated by SDS-PAGE using 10% or 12.5% Laemmli gel as described previously (90). For separation of the F-FecR protein and its derivatives, SDS-PAGE using a 15% Bis-Tris gel (107 mM bis-Tris (pH6.8 with HCl), 15% Acrylamide/Bis, 0.1% APS, 0.1% TEMED) and MES SDS running buffer (50 mM Tris, 50 mM MES, 1 mM EDTA, 0.1% (w/v) SDS) was used.

### Immunoblotting

Cells were grown at 30°C in M9-based medium with 20 μg/mL each of the 20 amino acids, 2 μg/mL thiamine and 0.4% glucose until mid-log phase. Total cellular proteins were precipitated with 5% trichloroacetic acid (TCA), washed with acetone and dissolved in SDS sample buffer. Immunoblotting was carried out essentially as described previously (90, 91). Proteins were separated by SDS-PAGE and electroblotted onto an Immobilon-P membrane filter (MilliporeSigma). Only when 15% bis-Tris gel was used for SDS-PAGE, a transferred membrane filter was dried at 37°C for 30 min and then hydrophilized with methanol. After blocking with BLOTTO (90), the filter was incubated with an appropriate antibody. For anti-RseP immunoblotting, anti-RseP antibodies were pre-incubated with whole-cell lysates of AD1840 (the Δ*rseA* Δ*rseP* Δ*degS* strain) at 4°C for 1 h to reduce a background as described previously (83). The filter was then washed, and incubated with goat anti-mouse or anti-rabbit IgG conjugated with horseradish peroxide (Bio-Rad). After washing of the filter, proteins that reacted with secondary antibodies were visualized using ECL or ECL Prime Western Blotting Detection Reagents (Cytiva) and Bio image analyzer LAS4000mini (Cytiva).

### Pulse-chase experiment

Cells were grown at 30°C in M9-based medium with 20 μg/mL each of the 18 amino acids (other than methionine and cysteine), 2 μg/mL thiamine and 0.4% glucose until early-log phase. In the experiments shown in Fig. 5 and Fig. S3B, the cells were induced with 1 mM IPTG and 1 mM cAMP for 10 min before pulse-labeling with [^35^S]methionine for 1 min and chase with an excess amount of unlabeled methionine for the indicated periods. In the experiments shown in Fig. 6A, B and Fig. S4A, cells were pulse-labelled with [^35^S]methionine for 1 min and chased with an excess amount of unlabeled methionine where 1.2 mM IPTG and 1.2 mM cAMP were added to the medium 5.5 min after the initiation of the chase. The proteins were precipitated with 5% TCA, washed with acetone and dissolved in 30 μL of 50 mM Tris-HCl (pH 8.1) containing 1 mM EDTA and 1% SDS by vigorous mixing for 30 min at room temperature followed by incubation at 37°C for 5 min. The samples were diluted 33-fold with 1 mL Triton buffer (50 mM Tris-HCl (pH 8.1), 0.1 mM EDTA, 2% Triton X-100, 150 mM NaCl). After clarification by centrifugation, the supernatants were mixed with anti-FLAG M2 Affinity Gel and Protein A Sepharose (Cytiva) and incubated at 4°C overnight. The antigen-antibody complexes were recovered by centrifugation, washed with Triton buffer and then with 10 mM Tris-HCl (pH 8.1) and dissolved in SDS sample buffer. The proteins were separated by SDS-PAGE and visualized and quantified using a phosphor imager (BAS5000) (Cytiva).

## Data availability

The MS raw data and analysis files have been deposited to the ProteomeXchange Consortium (http://proteomecentral.proteomexchange.org) via the jPOST partner repository (http://jpostdb.org) (92) with the data set identifier PXD023797.

## Acknowledgments

We thank Kazue Kanehara for constructing the plasmid pKK47. We thank Hiroyuki Mori, Ryoji Miyazaki, Eiji Ishii, and all the other lab members for stimulating discussion, and Michiyo Sano and Satomi Koshiba for secretarial support, and National BioResource Project (NBRP) –*E. coli* at National Institute of Genetics, Japan for strains. We also thank Editage for English language editing.

## Abbreviations

The abbreviations used are:

FL: full length
IMP: intramembrane protease
IPTG: isopropyl-*β*-D-thiogalactopyranoside
MBP: maltose binding protein
TM: transmembrane
WT: wild type

